# Characterisation of the bacterial microbiota of a landfill-contaminated confined aquifer undergoing intrinsic remediation

**DOI:** 10.1101/2020.05.28.120956

**Authors:** Daniel Abiriga, Andrew Jenkins, Kristian Alfsnes, Live S. Vestgarden, Harald Klempe

**Author notes:** Corresponding author. Department of Natural Sciences and Environmental Health, University of South-Eastern Norway, Gullbringvegen 36, NO-3800 Bø, Norway.

## Abstract

Literature on microbiome of landfill leachate-contaminated aquifers is scarce despite groundwater contaminations from landfills being common globally. In this study, a combination of microbiological techniques was applied to groundwater samples from an aquifer contaminated by a municipal landfill and undergoing intrinsic bioremediation. Groundwater samples were obtained from three multilevel sampling wells placed along the groundwater flow path in the contaminated aquifer and additionally from a background well located in a nearby uncontaminated aquifer. The samples were subjected to chemical analysis, microbial culturing and characterisation, cell counting by fluorescence microscopy and 16S rRNA metabarcoding. Good concordance was realised with the results from the different microbiological techniques. Samples from the uncontaminated aquifer had both lower cell density and lower microbial diversity compared to samples from the contaminated aquifer. Among the wells located in the contaminated aquifer, microbial diversity increased between the well closest to the landfill and the intermediate well, but was lower at the most distant well. The majority of the cultured microbes represented taxa frequently recovered from contaminated environments, with 47% belonging to taxa with previously documented bioremediation potential. Multivariate redundancy analysis showed that microbial composition was most similar in wells located closer to the landfill, although beta diversity analysis indicated a significant difference in microbial composition across the wells. Taken together with the results of cell counting, culture and metabarcoding, these findings illustrate the effect of landfill leachate on the microbial community and indicate that microbes are capable of hydrocarbon, sulphur, nitrogen, iron and manganese metabolism.

## Introduction

Groundwater is the main source of freshwater for drinking, agriculture and industry in many places globally (Mays and Scheibe, 2018; O’Connor et al., 2018), but faces serious pollution challenges (Brad et al., 2013; Pous et al., 2018; Röling et al., 2001) of which landfill is one. All over the world, landfills have served as the ultimate destination for municipal solid wastes (Reinhard et al., 1984), and continue to do so (Eggen et al., 2010). In Norway, there was little recycling of wastes until the late 1990s and most of the wastes from households and industries were deposited in municipal solid waste landfills with no provision for treatment or containment of the resultant leachate. Revdalen Landfill represents one such historic site. It was active from 1974 to 1997, leading to the contamination of Revdalen Aquifer. Pollutants of environmental concern such as heavy metals and polycyclic aromatic hydrocarbons have been detected in Revdalen Aquifer (Abiriga et al., 2020a).

Several strategies exist to reclaim contaminated aquifers. They are broadly categorised as artificial and natural. The former, which includes the conventional pump and treat (P&T) are faster, but require a major economic input for operation and maintenance (Hyldegaard et al., 2019). Natural attenuation such as *in situ* bioremediation on the other hand, offers inexpensive, eco-friendly yet efficient remedies (Logeshwaran et al., 2018; Nunes et al., 2013). In addition, unlike P&T, *in situ* bioremediation does not generate secondary wastes. It is therefore the most widely preferred option in *in situ* remediation of groundwater (O’Connor et al., 2018). However, *in situ* bioremediation, particularly intrinsic bioremediation is slow and the groundwater remains polluted for a long time, although it can be accelerated by amended bioremediation (Logeshwaran et al., 2018). Revdalen Aquifer is undergoing intrinsic bioremediation and the relevance of natural attenuation processes have been discussed previously (Abiriga et al., 2020a; b).

Traditionally, groundwater bioremediation has been demonstrated empirically by measuring geochemical parameters, with little use of microbial data (Mouser et al., 2005). Over the years, however, it has become apparent that studying microbial community composition in addition to geochemical measurements offers a more complete picture of bioremediation (Lu et al., 2012; Pilloni et al., 2019; Röling et al., 2001). In order to make inferences about bioremediation and effectively manage the processes, an understanding of the microorganisms responsible is necessary (Alfreider et al., 2002; Dlugonski, 2016; Köchling et al., 2015; Röling et al., 2000). A number of studies have been conducted on the microbiology of contaminated aquifers (Anantharaman et al., 2016; Cozzarelli et al., 2000; Harvey et al., 1984; Hug et al., 2015; Kleikemper et al., 2005; Ludvigsen et al., 1999; Röling et al., 2000; Tischer et al., 2012; Watanabe et al., 2002). Nonetheless, this area still requires more elucidation (He et al., 2018; Meckenstock et al., 2015). Bioremediation of hydrocarbon-polluted aquifers is well documented in the literature (Dojka et al., 1998; Harvey et al., 1984; Kleikemper et al., 2005; Pickup et al., 2001; Rooney-Varga et al., 1999; Tischer et al., 2012; Watanabe et al., 2002), but there is a dearth of studies on landfill leachate contaminations. The bias may reflect high-profile cases of hydrocarbon pollutions and the potential health hazard presented by the concomitant xenobiotics, which are often toxic, mutagenic and carcinogenic (Logeshwaran et al., 2018). Moreover, hydrocarbons, at least are, easily degraded in the environment and their compositions are less complex than effluents emanating from landfills. The complicated attenuation processes in landfill leachate impacted groundwater makes assessments of bioremediation processes more difficult (Christensen et al., 2000) and less attractive.

Studies on the microbiology of landfill-impacted aquifers (Albrechtsen et al., 1995; Lin et al., 2007; Ludvigsen et al., 1999; Mouser et al., 2005; Röling et al., 2000) have provided insights into the microbial profiles, but they employed low throughput methods that cannot capture the full microbial diversity. Fewer studies have applied high throughput sequence-based methods (Chen et al., 2017; Taş et al., 2018) and they have not specifically focused on microbial diversity. Thus, there is insufficient literature on the microbial diversity of landfill-perturbed aquifers. The present study aimed to describe the microbial diversity and composition of a landfill-leachate-contaminated confined aquifer using a combination of three microbiological techniques: culture and characterisation, cell counting by fluorescence microscopy and Illumina MiSeq 16S rRNA metabarcoding. To the best of our knowledge, no such multi-methodological study of landfill-perturbed subsurface microbiology has been reported.

## Materials and methods

### Study area and groundwater sampling wells

Revdalen Aquifer is a glaciofluvial deposit located in Vestfold and Telemark County in southeast Norway at coordinate 59°25′58.26″N and 9°06′1.53″E (Figure 1). In 1958, a kettle hole was adopted as a waste disposal site and through 1958-1974, the hole was directly filled up. Four successive landfill cells were later opened up in the area and were filled up between 1974 and 1996. Due to the leachate from the landfill, the aquifer, which was serving as a water source for the surrounding communities, became contaminated and water extraction was halted. Since the closure of the landfill in February 1997, the aquifer has been allowed to undergo monitored natural attenuation. Three multilevel monitoring wells were established along the groundwater flow direction to monitor the groundwater quality at 26 m (R1), 88 m (R2) and 324 m (R4) from the edge of the landfill. The wells are hereafter referred to proximal, intermediate and distal wells, respectively. In addition, a background well (R0) was established in a nearby aquifer for benchmarking the water quality. Additional information on the study site can be found elsewhere (Abiriga et al., 2020a; b).

**Figure 1.**
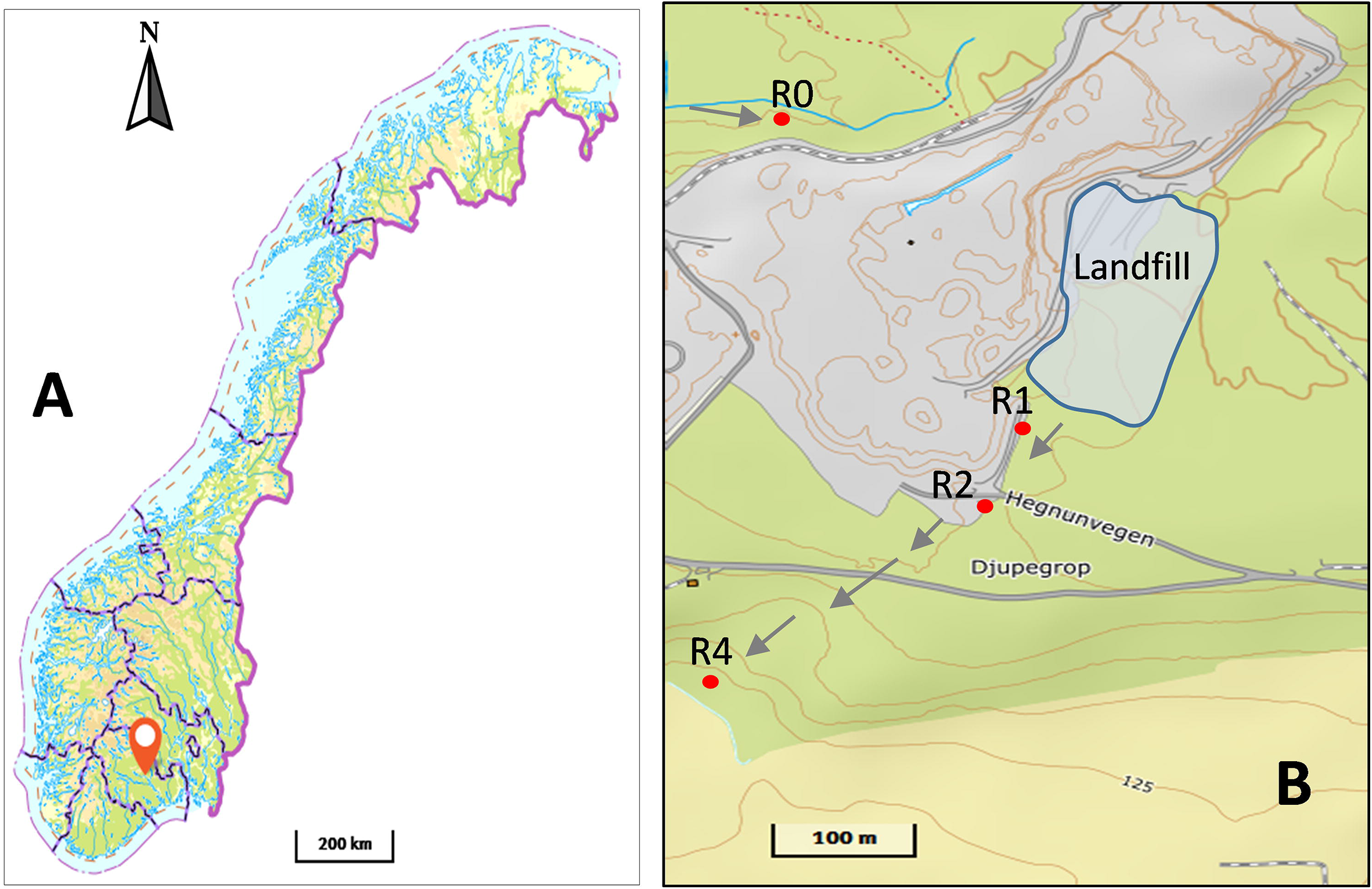
Location of the study site on map of Norway (**A**) and the location of landfill and sampling wells R0, R1, R2 and R4 (**B**) [Map source: Norwegian Geographical Survey, www.norgeskart.no, with permission]. R0 is the background well located in an uncontaminated, while wells R1, R2 and R4 are located in the contaminated aquifer placed at proximal, intermediate and distal positions, respectively. Arrows indicate groundwater flow direction. Green shading indicates woodland, yellow indicates farmland, and grey indicates industrial land, including the landfill and an adjacent active gravel/quarry pit. Contour interval 5 m.

### Groundwater sampling and chemical analysis

Groundwater sampling procedure has been described previously (Abiriga et al., 2020a) and was conducted as per International Organisation for Standardisation (ISO) guideline, ISO 5667-11(2009). Samples for this study were collected in spring and autumn 2018. For dissolved oxygen analysis, samples were collected in Winkler bottles, fixed onsite and protected from direct light. For other chemical analyses, samples were collected in 500 ml polyethylene bottles. pH and electrical conductivity were determined in the field using pH-110 meter (VWR International) and Elite CTS Tester (Thermo Scientific, Singapore), respectively. The samples were maintained at ≤4 □ using icepacks and a cooler box and transported to the laboratory at University of South-Eastern Norway. Samples for iron and manganese were filtered and acidified with nitric acid to pH ~2, while samples for total nitrogen were preserved by acidifying using sulphuric acid. All samples were stored at 4 □ until analysis. Norwegian Standards were followed for determination of dissolved oxygen (NS-ISO 5813), alkalinity (NS-EN ISO 9963-2), iron (NS 4773), and manganese (NS 4773). Major ions (ammonium, sodium, potassium, calcium, magnesium, chloride, nitrate and sulphate) were determined using Ion Chromatography DIONX ICS-1100 (Thermo Scientific, USA). Total nitrogen and total organic carbon (TOC) were determined using FIAlyzer-1000 (FIAlab, USA) and TOC Fusion (Teledyne Tekmar, USA), respectively.

### Microbial analyses

#### Sample handling

Groundwater samples were collected in sterile 350 ml PETE bottles (VWR, UK) and transported as described above. Upon arrival at the microbiology laboratory, 4.5 ml potions were fixed with 2.5% (final concentration) phosphate-buffered glutaraldehyde for microscopy. For 16S rRNA metabarcoding, 300 ml of water was filtered through 25 mm diameter 0.2 μm polycarbonate membrane filters. The filters were stored at −70 □ prior to DNA extraction. The remaining water was used for culturing heterotrophic bacteria. All the different sample handlings were conducted within 48 hours.

#### Direct microscopic count

1.9 ml of fixed groundwater samples were stained with 5 μg/ml 4□,6-diamidino-2-phenylindole (DAPI) (Kepner and Pratt, 1994). The stained cells were filtered onto 0.2 μm black polycarbonate Nuclepore membrane filters (Sigma-Aldrich, Germany), transferred onto microscope slides and overlaid with antifade mountant oil (Citifluor AF87, EMS, PA, USA). Cells were enumerated under X100 oil objective using Olympus IX70 fluorescence microscope (Tokyo, Japan). Ten fields were counted and the average count was used to estimate bacterial density using the formula: Bacteria (Cells/ml) = (*N* × *A*_*t*_)/(*V*_*f*_ × *A*_*g*_ × *d*), where *N* = average number of cells, *A*_*t*_ = effective area of the filter paper, *V*_*f*_ = volume of water sample filtered, *A*_*g*_ = area of the counting grid, and *d* = dilution factor (Kepner and Pratt, 1994). No observable cells were found in our blanks and therefore correcting for background noise due to contamination was not necessary.

#### Culturing and sequencing of heterotrophic bacteria

Preliminary assessment (data not shown) indicated dilutions >10^2^ could generate countable colonies and best growth occurred on half-strength tryptic soy agar. Thus, serial dilutions of 10^2^, 10^3^, and 10^4^ were prepared for each water sample and triplicates of 100 μl aliquots were spread on half-strength tryptic soy agar (20 g/l TSA and 7 g/l agar). Plates were incubated at 15 □ for >5 days. Aerobes were counted following incubation under aerobic condition. Anaerobes were enumerated in a parallel setup following incubation under anaerobic condition using GasPak with EZ Anaerobe Container System (BD, USA). Plates from two dilutions of each sample were counted and the average was reported as colony forming units (cfu) per ml.

Based on observable colony morphologies such as shape, elevation, margin, size, and colour, such as colonies covering the full diversity of colony morphologies were picked and purified by repeated streaking and incubation until pure cultures were obtained. These cultures were then observed by wet field microscopy at X1,000 (Olympus, CX22LEDRFS1, China) to observe motility and cell shape, followed by Gram staining and determination of oxidase and catalase activity (Csuros et al., 1999). Strains were stored at −70 □ in nutrient broth (Sigma, Switzerland) supplemented with 25% glycerol. The laboratory procedure for DNA extraction and sequencing of V3-V5 16S rRNA gene region of the pure isolates is provided as supplementary file (Method S1).

#### 16S rRNA metabarcoding

The frozen filters were retrieved and cut into two halves using sterile surgical blades and DNA was extracted from one-half using DNeasy PowerSoil Kit (Qiagen, Germany) following the manufacturer’s instruction. The DNA quantity and quality were checked using Qubit^®^ Flourometer 3.0 (Life Technologies, Malaysia) and 2% agarose gel electrophoresis, respectively. PCR and 16S rRNA gene library preparation (Fadrosh et al., 2014) for the samples were conducted at Norwegian Sequencing Centre (https://www.sequencing.uio.no). Both forward and reverse oligos included Illumina-specific adaptor sequence, a 12-nucleotide barcode sequence, a heterogeneity spacer and the primers 319F (5’-ACTCCTACGGGAGGCAGCAG-3’) and 805R (5’-GGACTACNVGGGTWTCTAAT-3’) for V3-V4 hypervariable 16S rRNA gene region. The pooled libraries were sequenced using Illumina MiSeq (600 cycles), applying the 300 bp paired-end protocol with 10% PhiX.

Sequence demultiplexing was conducted using a demultiplexer available at https://github.com/nsc-norway/triple_index-demultiplexing/tree/master/src. Barcodes and heterogeneity spacers were removed in the process, with no mismatches allowed in the barcode. Denoising (primer trimming, and removal of shorter sequences and chimeras), dereplication, merging, and clustering of sequences into amplicon sequencing variants were performed using DADA2 algorithm (Callahan et al., 2016) plug-in for QIIME2 version 2019.1.0 (Bolyen et al., 2019). Default parameters were implemented, with the exception of primer length (adjusting to 20 bp) and minimum sequence length of reads (adjusted to 280 bp). Taxonomic assignment was conducted using Naïve Bayes classifier algorithm trained on data from SILVA v.138 using QIIME2 version 2020.2.0.

##### Quality control

Quality control samples included in the sequencing run consisted of an elution buffer, extraction blanks and mock communities, which are described in more detail in the supplementary information (Method S2).

### Data availability statement

The 87 16S rRNA gene sequences have been deposited in GenBank under Accession Nos. MT348616-MT348702. The raw sequence reads from the metabarcoding have been deposited in Sequence Read Archive under BioProject PRJNA677875. The underlining data for this study were collected in 2018, with BioSamples SAMN16775936-SAMN16775961.

### Data analysis and statistics

All data analyses were conducted in R version 4.0.2 (R Core Team, 2020). To compare water chemistry between the background sample and the contaminated water samples, one-tailed Wilcox rank test was used. Comparison of water quality across the wells in the contaminated aquifer was performed using Kruskal-Wallis rank sum test. These tests were chosen as the majority of the variables showed non-normal distribution. One-tailed paired *t*-test was used for both within-sample and overall comparison between plate and microscopic counts. Similarly, comparison between aerobic and anaerobic counts was also done using one-tailed paired *t*-test. One-way ANOVA was conducted to test for differences in aerobic and microscopic counts across the wells and a post-hoc Tukey’s Honest Significant Difference for the pairwise comparisons. All the parametric tests were performed on log-transformed data. Normality of data distribution was assessed graphically using histograms, boxplots and by Shapiro-Wilk normality test. Statistical significance was inferred at alpha = 0.05.

Multivariate analyses were performed using package vegan in R (Oksanen et al., 2019). Prior to redundancy analysis (RDA), the OTU abundance data was pre-transformed using fourth-root transformation to reduce asymmetry of the data distribution before standardising it using Hellinger standardisation (Legendre and Gallagher, 2001). Also, because the groundwater physicochemical variables are dimensionally heterogeneous, the chemistry data was scaled prior to RDA ordination. The difference in the community profiles of the sample groups in the RDA triplot was tested using permutational multivariate analysis of variance on Euclidean distance. The homogeneity of group dispersion was assessed beforehand using function *betadisper*. Principal component analysis (PCA) was applied on square-root transformed and Hellinger-standardised culture-based microbial data and physicochemical data fitted using command *envfit* in package vegan. Taxonomic-to-phenotypic mapping was implemented using METAGENassist (Arndt et al., 2012).

## Results

### Groundwater chemistry

Most of the solutes in the groundwater were present at levels significantly above that of the background well (Table 1). Kruskal-Wallis test showed significant difference across the wells in the contaminated aquifer for eight of the fifteen variables measured (Table S1). While the majority of the variables showed decrease in concentration along the groundwater flow path, a few i.e. dissolved oxygen, magnesium, manganese and iron showed increase in concentration downgradient.

**Table 1.**
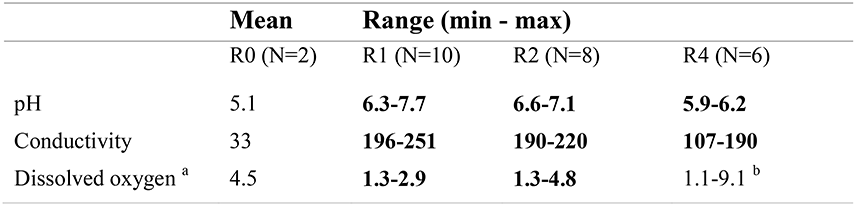
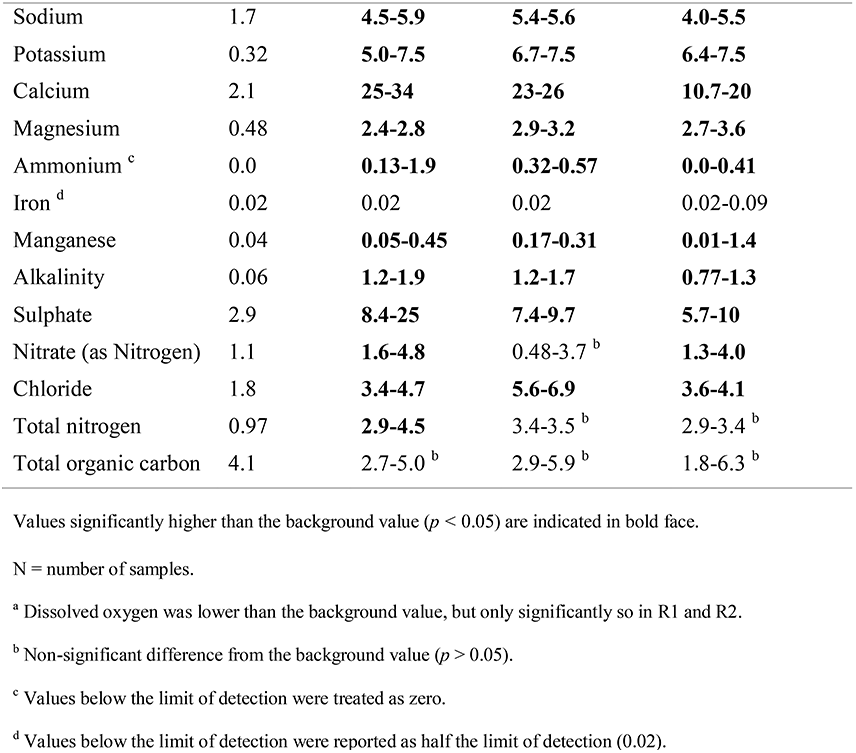
Characteristics of the groundwater chemistry measured in spring and autumn 2018. Values from the background well (R0) were used as a benchmark against which those in the proximal (R1), intermediate (R2) and distal (R4) were compared. All units are in mg/l, except for pH (pH units), conductivity (μS/cm) and alkalinity (mM).

|  | Mean | Range (min - max) |  |  |
| --- | --- | --- | --- | --- |
|  | R0 (N=2) | R1 (N=10) | R2 (N=8) | R4 (N=6) |
| pH | 5.1 | **6.3-7.7** | **6.6-7.1** | **5.9-6.2** |
| Conductivity | 33 | **196-251** | **190-220** | **107-190** |
| Dissolved oxygen <sup>a</sup> | 4.5 | **1.3-2.9** | **1.3-4.8** | 1.1-9.1 <sup>b</sup> |
| Sodium | 1.7 | **4.5-5.9** | **5.4-5.6** | **4.0-5.5** |
| Potassium | 0.32 | **5.0-7.5** | **6.7-7.5** | **6.4-7.5** |
| Calcium | 2.1 | **25-34** | **23-26** | **10.7-20** |
| Magnesium | 0.48 | **2.4-2.8** | **2.9-3.2** | **2.7-3.6** |
| Ammonium <sup>c</sup> | 0.0 | **0.13-1.9** | **0.32-0.57** | **0.0-0.41** |
| Iron <sup>d</sup> | 0.02 | 0.02 | 0.02 | 0.02-0.09 |
| Manganese | 0.04 | **0.05-0.45** | **0.17-0.31** | **0.01-1.4** |
| Alkalinity | 0.06 | **1.2-1.9** | **1.2-1.7** | **0.77-1.3** |
| Sulphate | 2.9 | **8.4-25** | **7.4-9.7** | **5.7-10** |
| Nitrate (as Nitrogen) | 1.1 | **1.6-4.8** | 0.48-3.7 <sup>b</sup> | **1.3-4.0** |
| Chloride | 1.8 | **3.4-4.7** | **5.6-6.9** | **3.6-4.1** |
| Total nitrogen | 0.97 | **2.9-4.5** | 3.4-3.5 <sup>b</sup> | 2.9-3.4 <sup>b</sup> |
| Total organic carbon | 4.1 | 2.7-5.0 <sup>b</sup> | 2.9-5.9 <sup>b</sup> | 1.8-6.3 <sup>b</sup> |

Values significantly higher than the background value ( $p < 0.05$ ) are indicated in bold face.
N = number of samples.
<sup>a</sup> Dissolved oxygen was lower than the background value, but only significantly so in R1 and R2.
<sup>b</sup> Non-significant difference from the background value ( $p > 0.05$ ).
<sup>c</sup> Values below the limit of detection were treated as zero.
<sup>d</sup> Values below the limit of detection were reported as half the limit of detection (0.02).

### Groundwater microbiology

#### Viable plate count

The cell density estimate from the plate count was in the range 1×10^2^ - 3.2×10^5^ cfu/ml (aerobic) and 0 - 2.4×10^5^ cfu/ml (anaerobic). Despite the comparable maximum counts from the two growth conditions, the aerobic plate count was significantly higher than the anaerobic plate counts (*t* = 3.63, *df* = 37, *p* = 0.0004***). The same strains dominated under both aerobic and anaerobic conditions and isolates recovered under anaerobic condition were all facultative anaerobe. Nonetheless, some strains were isolated only under anaerobic condition. Wells in the contamination plume had higher plate counts than the background well and the distal well had lower plate counts compared to the intermediate and proximal wells (Figure 2). The count was significantly different across the wells (*F* = 3.09, *df* = 3, *p* = 0.0357*), but pairwise comparisons between the wells did not give significant results.

**Figure 2.**
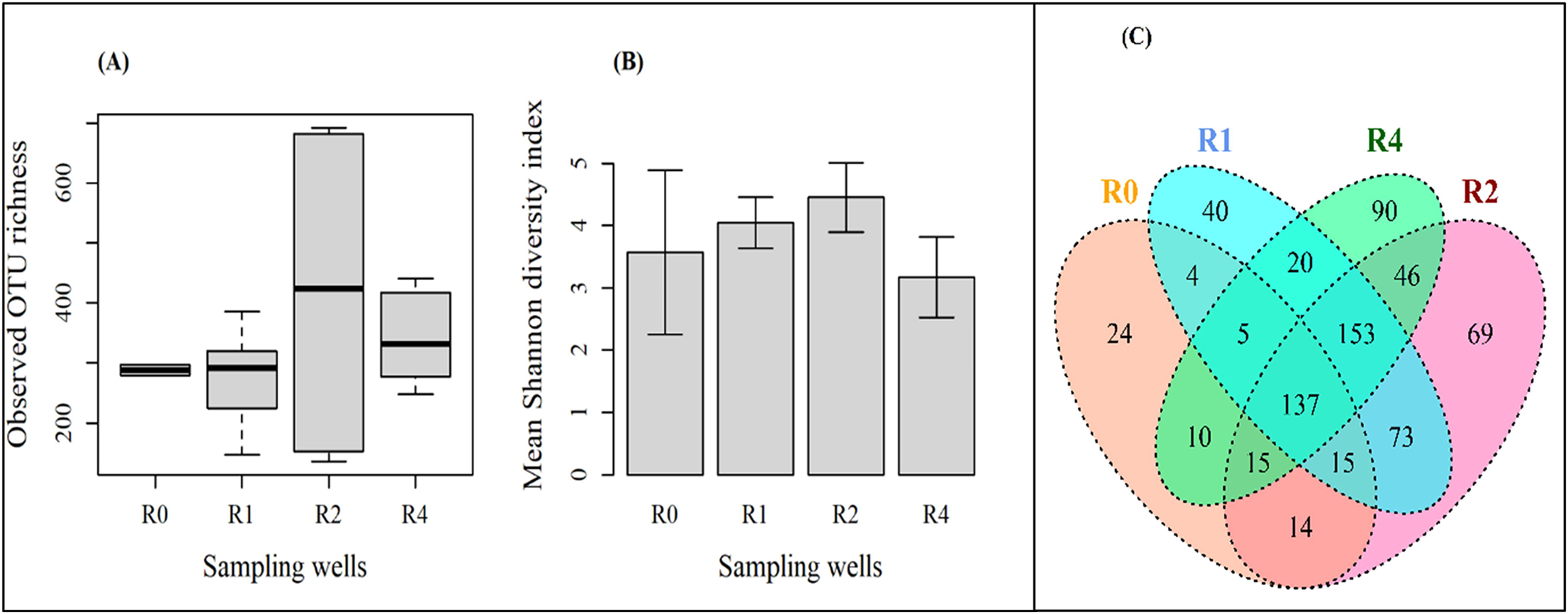
Bacterial cell density in the groundwater samples estimated from direct microscopic count (MC) and aerobic plate count (PC); error bars are mean + standard error (n = 4). R0 is the background well located in an aquifer upstream of the landfill. R101-R105, R201-R204 and R401-R403 are the multilevel in R1, R2 and R4, and are the wells located in the contaminated aquifer placed along the groundwater flow direction at the proximal, intermediate and distal positions, respectively. Blue asterisk (□) indicate significant difference.

#### Direct fluorescence microscopy

Microscopic counts were in the range 7×10^3^ - 3.5×10^5^ cells/ml. Cell morphologies observed under fluorescence microscopy included small rods, long large rods, coccobacilli and vibrios. Small rods were most frequently encountered. Water samples from the proximal and intermediate wells were dominated by short rods, vibrios and elongated narrow rods, some occurring in chains. The distal well on the other hand, did not exhibit predominance of any specific cell types, an observation that agrees with the more diverse colony morphology (higher richness) observed in samples from this well (see below). The total cell count was not significantly different across the four wells (*F* = 1.44, *df* = 3, *p* = 0.243), nor were the pairwise comparisons.

The distal well, which showed very low plate counts, gave a relatively higher microscopic count. Within-sample comparison indicates that microscopic counts were in all but one sample, higher than plate counts, but only significantly so in four of the thirteen samples: R0, R102, R104 and R204 (Figure 2). Overall, the microscopic count was higher than the plate count (*t* = 6.94, *df* = 51, *p* < 0.05***) (Figure S1).

#### Identification of isolates

Small subunit rRNA gene sequences of the pure isolates revealed higher species richness in the wells located in the contamination plume than in the background well (Figure S2). The distal well had the highest species richness. Eighty-seven taxa belonging to forty-six genera were identified (Figure S2, Table S2), *Pseudomonas* being the most abundant taxon (Figure S4). Of the eighty-seven taxa, seven were found only in the background aquifer, seventy-seven found only in the contaminated aquifer and three were found in both. These three were *Rhodoferax ferrireducens*, *Pseudomonas* sp. (2) and *Rugamonas* sp. detected in both the background and distal wells (Table S2). As shown in Figure 3, twenty-eight taxa were unique to the distal well. No single taxon was found in all the four wells, although two genera (*Pseudomonas* and *Janthinobacterium*) were common. Among the wells located in the contamination plume, only two species i.e. *Pseudomonas veronii* and *Rhodococcus degradans* were found in all of them. The proximal-intermediate was the most similar pair of wells, with ten species in common. PCA ordination further depicts this similarity (Figure S5).

**Figure 3.**
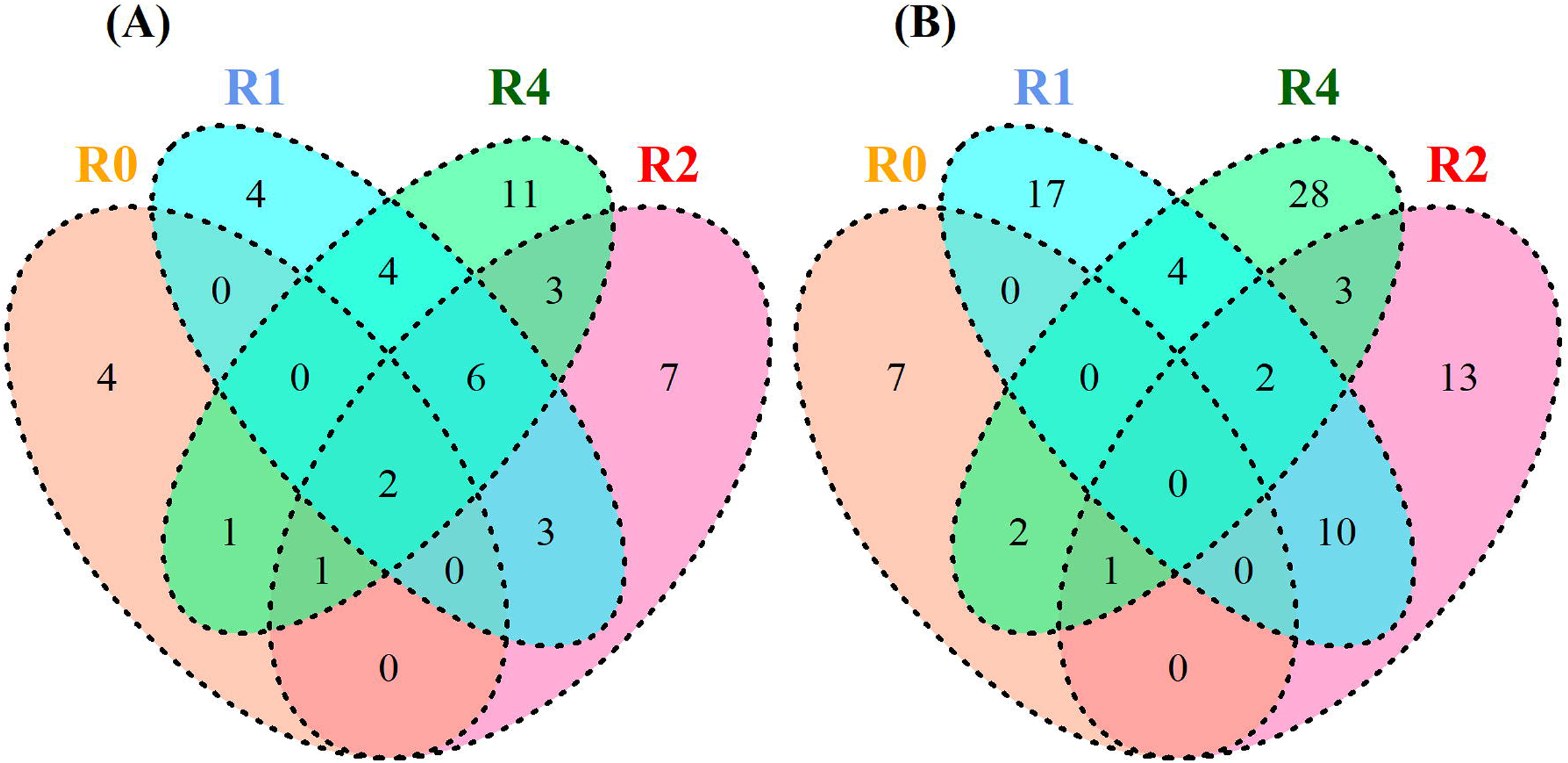
Venn diagram showing relationships among the groundwater sampling wells at genus-level classification **(A)** and species-level classification **(B)**. R0 is the background well located in an aquifer upstream of the landfill. R1, R2 and R4 are wells located in the contaminated aquifer placed along the groundwater flow direction at the proximal, intermediate and distal positions, respectively.

#### 16S rRNA metabarcoding

A total of 1496 OTUs were identified by metabarcoding. 98.73% were bacteria; the rest were archaea (1.2%) or unclassified (0.07%). These were classified to 57 microbial phyla, with *Proteobacteria* (31%), *Patescibacteria* (9.2%), *Bacteroidetes* (7.8%), *Actinobacteria* (6.5%), *Chloroflexi* (6.1%), *Acidobacteria* (5.5%), *Verrucomicrobia* (5.4%) and *Firmicutes* (3.5%) dominating in all the samples.

For the sake of simplicity and in order to target taxa likely to be stable members of the microbial community, only OTUs comprising at least 2% of at least one sample are considered here. This resulted in only 68 genera being represented (Figure 4), of these, 15 had unresolved taxonomic classifications and 17 were OTUs of uncultured microbes. Twelve of the thirty-six (33%) phylogenetically well-resolved genera were among those isolated through the culture-based approach. These were *Brevundimonas*, *Flavobacterium*, *Janthinobacterium*, *Pedobacter*, *Phyllobacterium*, *Polaromonas*, *Pseudomonas*, *Rhodanobacter*, *Rhodoferax*, *Sphingomonas*, *Stenotrophomonas* and *Undibacterium*. These genera also showed greater abundance in the PCA ordination for the cultured taxa (Figure S5). The 12 genera accounted for 38 of the total cultures isolated (Figure S6).

**Figure 4.**
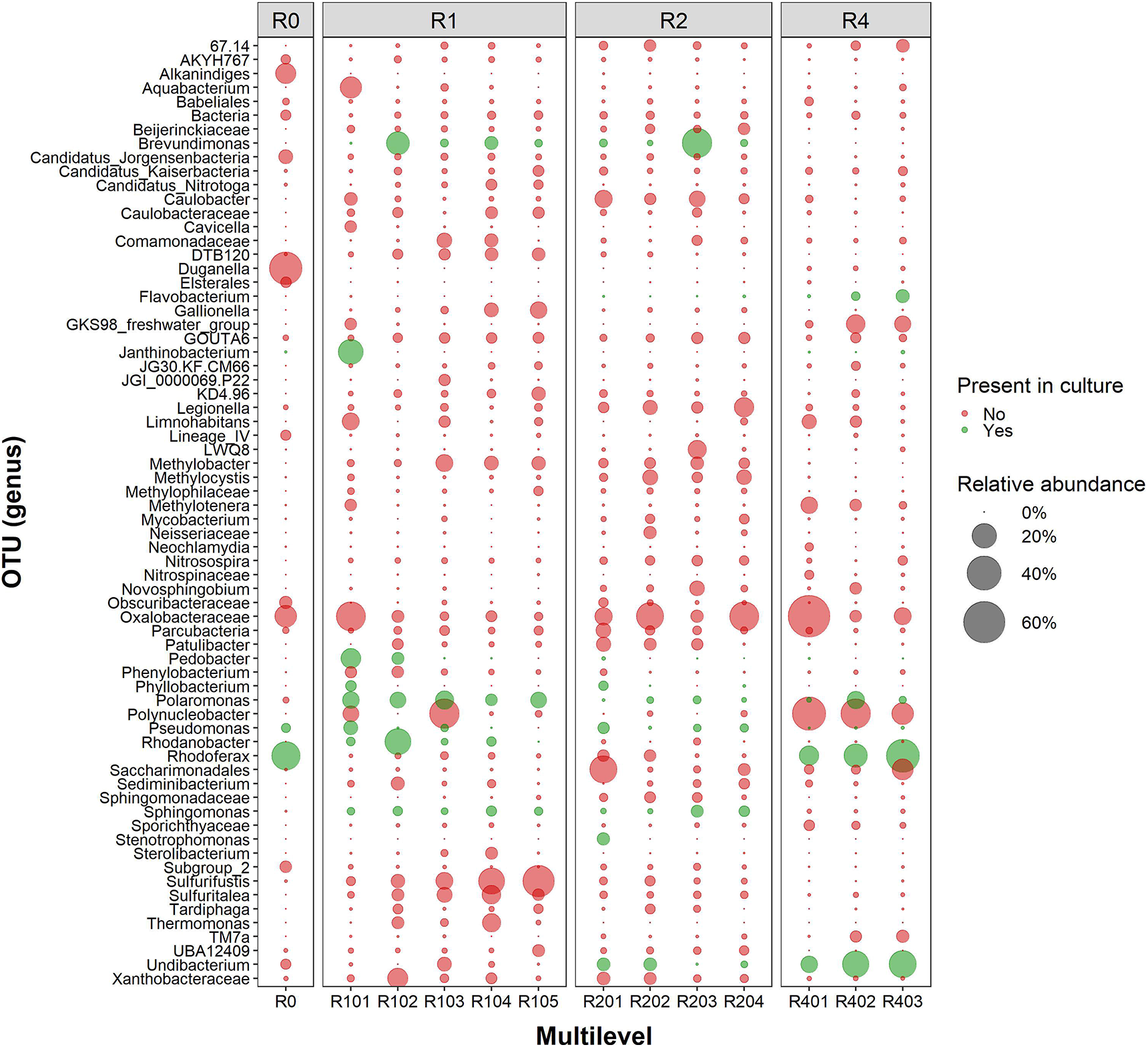
OTUs greater than 2% of the total reads in at least one of the samples. R0 is the background well located in an aquifer upstream of the landfill. R101-R105, R201-R204 and R401-R403 are the multilevel in R1, R2 and R4, and are the wells located in the contaminated aquifer placed along the groundwater flow direction at the proximal, intermediate and distal positions, respectively. For clarity, the relative abundances for spring and autumn were merged.

Shannon diversity measure was applied to the OTU data to infer species richness and diversity. Microbial diversity was greatest in the intermediate well, followed by the proximal well and was lowest in the distal well (Figure 5 and Table S3). Similarly, the evenness estimate followed the same trend, indicating that the distal well was dominated by few taxa, while the intermediate well had fairly even representation of microbial taxa present at the site.

**Figure 5.**
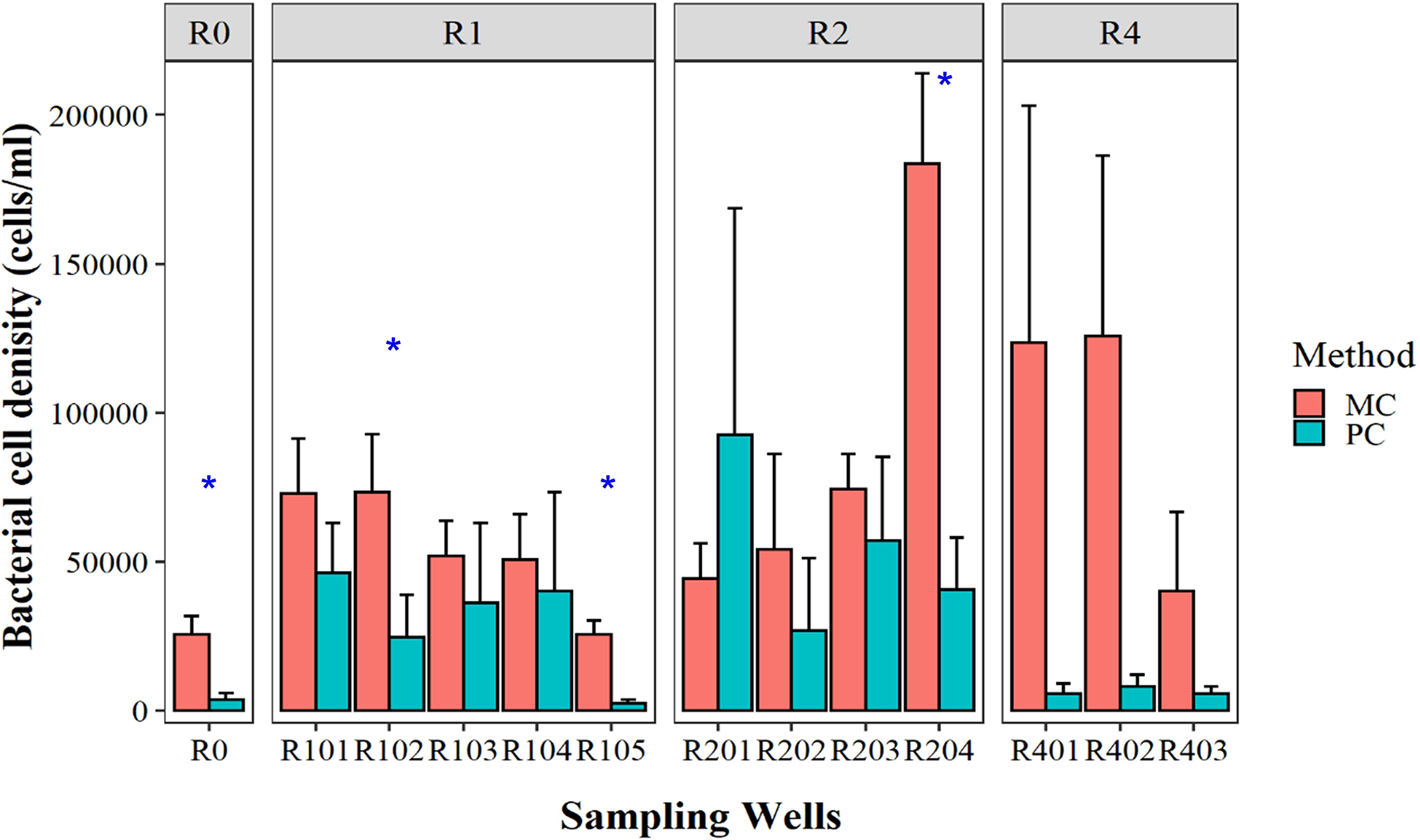
OTU richness **(A)**, Shannon diversity index **(B)** (error bars are ±SD) and species count at genus level **(C)** recorded in the sampling wells R0, R1, R2 and R4. R0 is the background well located in an aquifer upstream of the landfill. R1, R2 and R4 are the wells located in the contaminated aquifer placed along the groundwater flow direction at the proximal, intermediate and distal positions, respectively.

However, both the highest (692) and lowest (136) OTU richness was recorded at the different levels in the intermediate well. This resulted in a larger spread of OTU richness in the intermediate well (Figure 5). A count of species (at genera level) (Figure 5) indicates that a core of 137 genera were common to all the sampling wells, while a further 153 genera were common to wells of the contaminated aquifer. The number of unique genera varied between 24 in the background well and 90 in the distal well. The number of genera shared between just two wells varied from 4 (background/proximal) to 73 (distal/intermediate).

Multivariate analysis using RDA showed that microbial composition varied spatially, with the intermediate and proximal wells clustering next to each other as was observed with the culture-based method (Figure 6 and Figure S5). PERMANOVA indicates that the microbial compositions of the wells are significantly different (*F* = 4.58, *df* = 3, *p* = 0.001***). Three canonical axes were significant (*p* = 0.001, 999 permutations) and the model explained 34.9% (adjusted R^2^) of the variation in microbial composition. The proximal and the intermediate wells correlated positively with RDA1, while the distal and background wells correlated negatively with RDA1. The second axis (RDA2) separated mainly the background well from the rest of the wells located in the contaminated aquifer.

**Figure 6.**
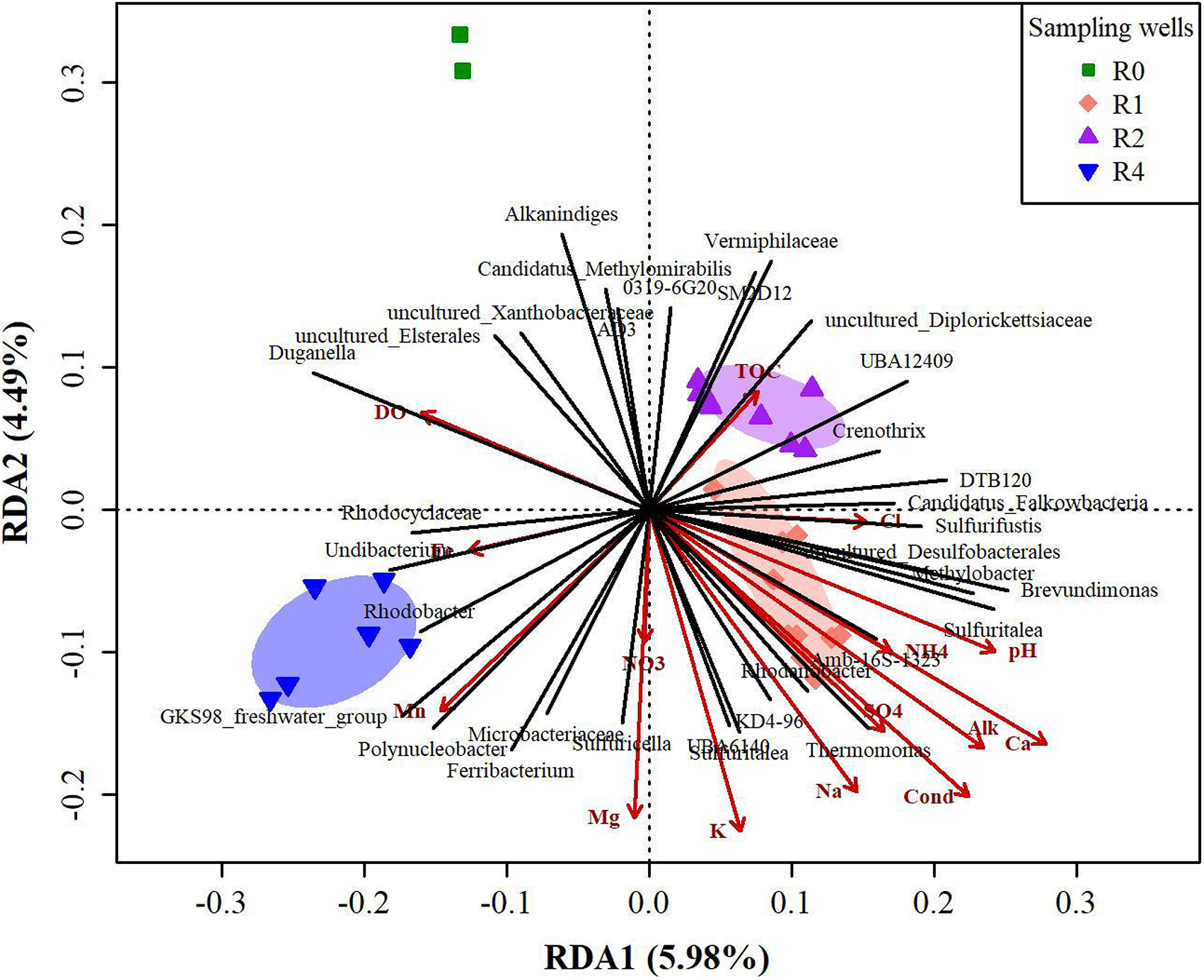
Redundancy analysis (RDA) of microbial composition among the sampling wells; background well (R0), proximal well (R1), intermediate well (R2) and distal well (R4). RDA was performed on Hellinger standardised and fourth-root transformed microbial composition data constrained by environmental variables conditional on season. For clarity of readability, only OTUs with prominent vector length are shown.

The groundwater physicochemical parameters have shown two major gradients: those that showed higher levels towards the proximal well and those that showed higher levels towards the distal well. The former included pH, ammonium, calcium, alkalinity, sulphate, conductivity, sodium, chloride and potassium, which have correlated negatively with RDA2. The second gradient was driven by dissolved oxygen, magnesium, iron and manganese, which have likewise correlated negatively with RDA2, except dissolved oxygen that correlated positively with RDA2. Nitrate and magnesium have shown moderate gradient between the proximal and distal wells, while TOC showed higher levels towards the intermediate well.

The prominent OTUs in the intermediate and background wells were dominated by uncultured lineages. On the other hand, the abundant OTUs in the distal and proximal wells composed mainly culturable taxa, although those in the proximal well seem to constitute those requiring special growth conditions. Some of the OTUs that showed prominent vectors in the RDA analysis were also among the abundant (>2%) OTUs. They include *Alkanindiges*, *Duganella*, *Undibacterium*, GKS98_freshwater_group, *Sulfuritalea*, *Sulfurifustis*, *Thermomonas*, *Rhodanobacter*, *Brevundimonas*, *Methylobacter*, DTB120 and UBA12409.

#### Predicted phenotypic functions

Taxonomic-to-phenotypic mapping using METAGENassist (Figure 7) predicted 15 potential energy source phenotypes, predominantly phototroph, heterotroph, autotroph and organotroph, methanotroph, methylotroph and oligotrophy in varying abundance varied from well to well. Five groups were present at a relative abundance >0.2. These were phototroph in the background and distal wells, heterotroph in the background and proximal wells, organotroph in the proximal well, and autotroph and methanotroph in the intermediate well. Phototroph and heterotroph accounted for >50% of the total abundance in the background and distal wells. Overall, the background well was dominated by two energy source-types, the proximal and intermediate wells by seven and six, respectively, and the distal well by four.

**Figure 7.**
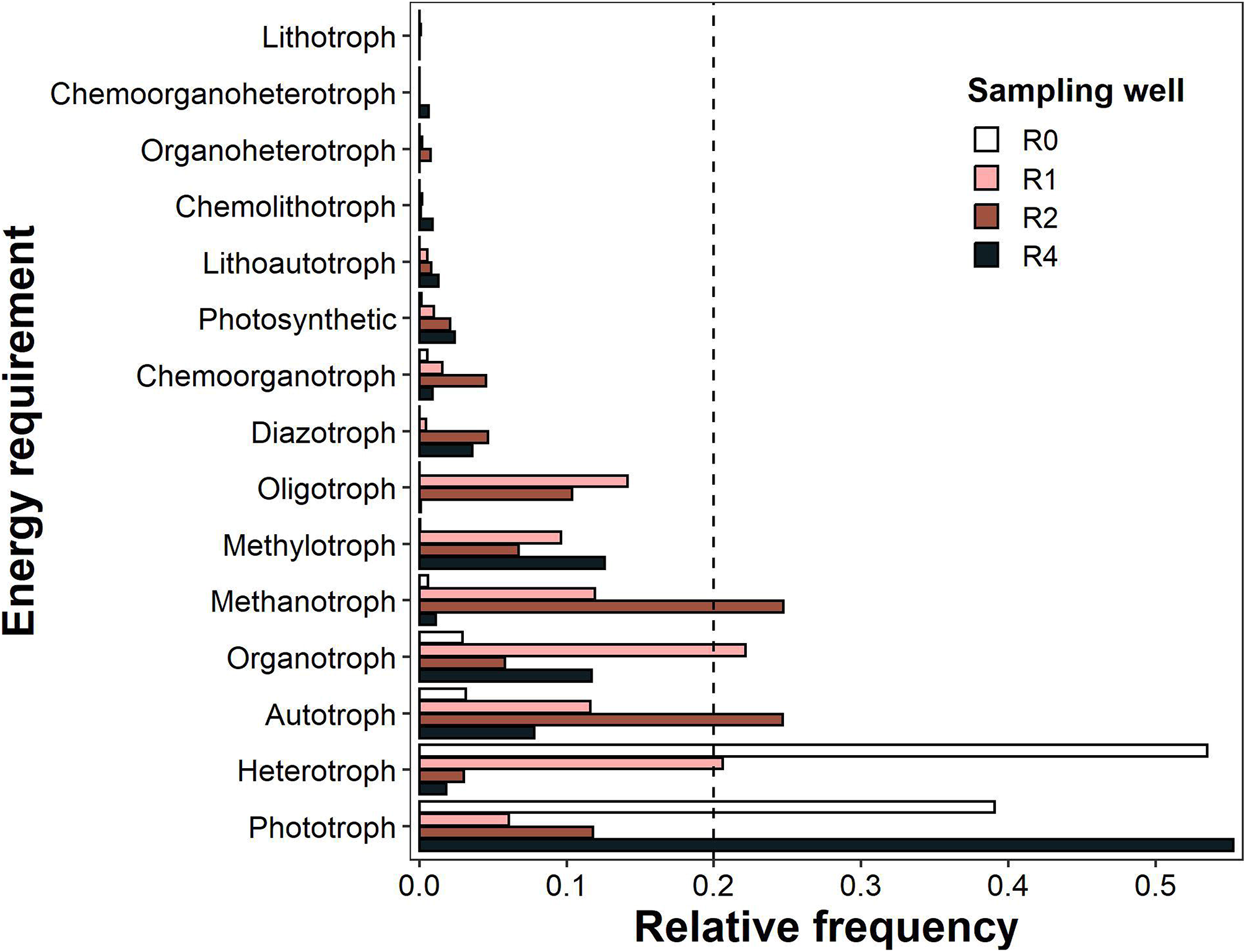
Predicted energy source requirements of the OTUs in R0, R1, R2 and R4. R0 is the background well located in a nearby uncontaminated aquifer, while R1, R2 and R4 are the wells located in the contaminated aquifer placed at the proximal, intermediate and distal positions from the landfill. The vertical dashed line depicts 20% abundance.

Twenty-seven different potential metabolic profiles were predicted (Figure 8). Degraders of both inorganic and organic compounds were predicted. Organic compound transformers were however, in lower abundances than inorganic compound metabolisers. Microbes involved in sulphur and nitrogen transformations formed the dominant groups. Generally, the metabolic profiles varied in abundance from well to well. With the exception of iron-reducers, streptomycin producers, and chitin, atrazine, chlorophenol, lignin and propionate degraders, most of the metabolic profiles showed greater abundance in the wells located in the contaminated aquifer.

**Figure 8.**
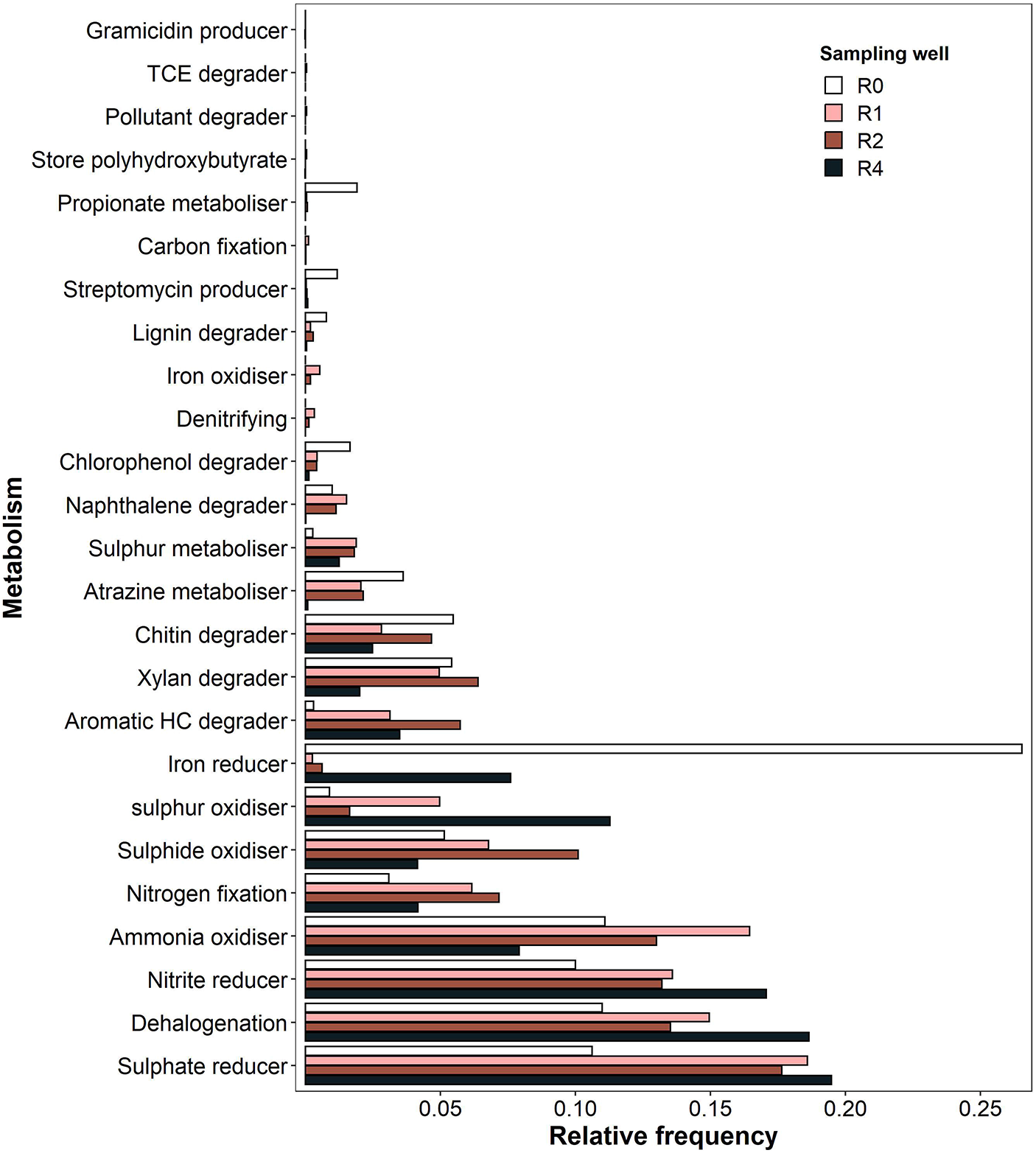
Predicted metabolic potentials of the OTUs in R0, R1, R2 and R4. R0 is the background well located in a nearby uncontaminated aquifer, while R1, R2 and R4 are the wells located in the contaminated aquifer placed at the proximal, intermediate and distal positions from the landfill.

## Discussion

### Cell density and microbial diversity

Overall, cell density estimate from microscopic count was significantly higher than plate count, which is consistent with the literature (Gregorich and Carter, 2007; Muyzer and Smalla, 1998) and expected from theoretical considerations. However, within the wells located in the contaminated aquifer, this difference was seldom greater than a factor of two in most cases, and only in three of the twelve samples was the difference significant. This suggests that in most samples, a large proportion of the microbial population is culturable. Presumably, the compounds in the landfill leachate favoured the growth of culturable heterotrophic microorganisms. The physiological profiles of microbial communities in aquifers receiving a landfill leachate is expected to be due to culturable bacteria (Röling et al., 2000). However, the metabarcoding data conflict with these findings, the highest proportion of uncultured OTUs being found in the intermediate well, rather than in the distal well as indicated by comparison of microscopic and plate counts.

The background aquifer had low solutes levels, indicating that the aquifer is nutrient-poor. Based on the culture method, both cell density and the microbial diversity were low, although comparison of microscopic and plate counts suggests that a fairly large proportion of the population is non-culturable. This is consistent with the metabarcoding data, in which the microbiome of the background aquifer was dominated by uncultured taxa and only a few culturable taxa such as *Duganella*, *Rhodoferxa* and *Alkanindiges* were abundant. Similarly, the Shannon diversity index (3.57 ± 1.32) was lower than the overall diversity index recorded in the contaminated aquifer (3.96 ± 0.71). Further, the RDA ordination indicates the background well is compositionally very different from the contaminated aquifer. Similar differences in microbial communities between contaminated and uncontaminated groundwater have been reported (Brad et al., 2013; Brad et al., 2008; Mouser et al., 2005).

The contaminated aquifer, on the other hand, is relatively solute-rich and supports a denser and more diverse microbial population; a scenario likely to occur where a landfill leaches easily degradable organic matter (Röling et al., 2000). Although the concentrations of solutes have decreased greatly compared to previous data (Abiriga et al., 2020a), they are evidently still sufficient to maintain a distinctive bacterial community; a previous study (Herzyk et al., 2017) indicates that physicochemical parameters return to normal more quickly than the microbial community. In line with this, data from culturing, fluorescence microscopy and metabarcoding showed higher species and cell density, diversity and species richness in the contaminated aquifer. The highest species richness and Shannon entropy were recorded in the intermediate well (Figure 5 and Table S3), an observation that agrees with the culture and microscopic methods, where the highest cell and plate counts were recorded in the intermediate well (Figure 2). On the other hand, moderate and lowest species richness and diversity were recorded in the proximal and distal well, respectively (Figure 5). This suggests an existence of ecological gradient along the groundwater flow path, in which the proximal and intermediate wells are expected to have a high resemblance, as they are spatially close to each other. This is consistent with both the culture and metabarcoding data (Figure 3 and Figure 5C), where the highest shared species and genera was recorded between the proximal and intermediate wells. Further, these wells clustered next to one another (Figure 6), suggesting that they are relatively similar and are thus made up of fairly similar microbial communities. The distal well is therefore considered to be ecologically more different from the upstream wells, which is consistent with the observations that the highest unique species (28) (Figure 3B) and genera (90) (Figure 5C) were found in the distal well.

The concentration of some of the groundwater physicochemical variables was found to decrease with distance from the landfill (Table 1), and this was accompanied by an increase in microbial diversity (Figure 5 and Table S3) at least from the proximal to intermediate wells. This may be analogous to increase in diversity and community stability as leachate becomes less contaminated over time (Köchling et al., 2015). Thus, close to the landfill, only species resistant to the toxic effects of the leachate are probably able to survive and grow, while as toxic pollutants become attenuated through biotic and abiotic processes, it allows the growth of more sensitive species. A previous study from Norman Landfill in the United States has shown that microbial gene diversity varied with distance from the landfill (Lu et al., 2012). In either study (both the previous and the present), the observed changes in diversity as a function of distance indicates the significance of attenuation mechanism *in situ* that shapes the microbial composition and function. Such spatial variation may enhance biodegradation as pollutants migrate from one region with a given microbial composition to another of a different microbial composition (Brad et al., 2013; Brad et al., 2008; Mouser et al., 2005; Röling et al., 2000).

### Microbial compositions and environmental significance

The culture-based analysis identified four microbial phyla (Figure S3), while the metabarcoding analysis recorded fifty-seven phyla of which only eight occurred in greater (>3.5%) abundances. Of these eight phyla, *Proteobacteria*, *Bacteroidetes*, *Actinobacteria* and *Firmicutes* were represented among the cultured isolates, while *Patescibacteria*, *Chloroflexi*, *Acidobacteria* and *Verrucomicrobia* were only present in the metabarcoding data. These latter phyla are largely represented by poorly-described and/or uncultured species, which limits inference of their physiological capabilities, and are not discussed further. *Proteobacteria* was the most abundant phyla, which accounted for 31% and 55.2% of the total microbiome in the metabarcoding and culture data, respectively. *Bacteroidetes* and *Actinobacteria* were present in fairly comparable proportions, while *Firmicutes* was the least abundant of the eight phyla. The ecological/environmental significance of the abundant (>2%) OTUs and pure isolates are discussed based on bibliography and phenotypic mapping from METAGENassist (Arndt et al., 2012).

#### Hydrocarbon metabolism

Potential hydrocarbon degraders have been identified by both metabarcoding and culture methods. The culture data revealed the presence of bacteria capable of degrading a wide range of both complex and simple hydrocarbons. Among them were degraders of polychlorinated biphenyls (PCB), diesel oil, crude oil, 4-chlorophenol, pentachlorophenol, trichloroethene, and polycyclic aromatic hydrocarbons (PAHs) such as phenanthrene, naphthalene and Benzo[a]pyrene (Table S4). Whether the presence of these bacteria was due to the organic compounds cannot be answered by the present study, but congeners of PAH have been detected in the aquifer water (Abiriga et al., 2020a). From the metabarcoding data, potential hydrocarbon metabolisms include antipyrine/chloridazon degradation by *Phenylobacterium* (Lingens et al., 1985; Oh and Roh, 2012), alkane/alkyl degradation by *Parvibaculum* and *Alkanindiges* (Bogan et al., 2003; Lai et al., 2011; Schleheck et al., 2004), ibuprofen degradation by *Patulibacter medicamentivorans* (Almeida et al., 2013) and oil degradation by *Aquabacterium* (*A. olei*) (Jeong and Kim, 2015). Other carbon metabolisms include a single-carbon metabolism by *Methylobacter* (Bowman et al., 1993), *Methylotenera* (Kalyuzhnaya et al., 2006) and *Methylocystis*. Energy source prediction showed the presence of methylotrophs and methanotrophs in high proportions in the contaminated wells (R1-R4) (Figure 7), which may suggest the presence of methane in the aquifer. The landfill has now matured (manuscript in preparation), and is supposedly in transition from methanogenic to aerobic stage, i.e. from phase III to phase IV of landfill stabilisation (Kjeldsen et al., 2002), but there could still be active methanogenesis. A wide range of hydrocarbon metabolisms have been predicted through the phenotypic mapping (Figure 8). Although the TOC in the groundwater was low with a weak gradient along groundwater flow path, it could still supply carbon sources to the aquifer microbiome. Considering the age of the landfill, the TOC should be predominantly recalcitrant fractions (Kulikowska and Klimiuk, 2008) and this makes it difficult to degrade (Appelo and Postma, 2005), leading to non-significant difference across the wells.

#### Sulphur, nitrogen and iron transformations

From the metabarcoding data, sulphur cyclers included sulphur/sulphide oxidisers *Sulfuritalea* (Kojima and Fukui, 2011), *Sulfurifustis* (Kojima et al., 2015) (Figure 4), and *Sulfuricella* (Kojima and Fukui, 2010) and *Rhodobacter* (Imhoff et al., 1984) (Figure 6). This is in agreement with the metabolic function prediction, which revealed presence of sulphur/sulphide oxidisers. Despite the implied abundance of sulphur/sulphide oxidisers, sulphate showed a weak gradient with a non-significant difference across the wells (Table S1), suggesting that the aquifer contains a significant sulphate sink, such as re-reduction in local anaerobic pockets.

Both nitrate reducers and ammonia oxidisers were identified. Nitrate reducers included 29 isolates (Table S5), and genera *Caviccella*, *Sterolibacterium*, *Aquabacterium* and *Novosphingobium* from the metabarcoding data. *Sulfuricella*, which at present is a monotypic (*S. denitrificans*), correlated positively with nitrate and could therefore, be involved in denitrification. Among the nitrogen oxidisers were the ammonia oxidisers *Nitrosospira* (Watson, 1971) and nitrite oxidisers *Nitrospinaceae* (Lücker and Daims, 2014). Taxonomic-to-phenotypic mapping (Figure 8) indicates the presence of ammonia oxidisers and nitrate/nitrite reducers in greater abundances. Both ammonium and nitrate were detectable in the groundwater, although nitrate was present in higher levels than ammonium. The presence of both ions may suggest complete cycling of nitrogen, although the reduction reaction might be limited due to low organic matter. The lack of significant difference in nitrate across the wells suggests a balance between reductive and oxidative nitrate metabolism.

Iron metabolisers include *Gallionella*, a genus known for centuries to clog well screens (Chapelle, 2001) by oxidising ferrous iron at an ecotone between reducing and oxidising environments. Genus *Rhodoferax*, which the culture-based analysis showed to be represented by *R. ferrireducens*, together with genus *Ferribacterium*, are both iron-reducing bacteria; a role that counteracts that played by *Gallionella*. In addition, *R. ferrireducens* is known to carry out metabolism of manganese and oxygen (Finneran et al., 2003). Although the levels of iron was low across the wells, the distal well where *Rhodoferax* and *Ferribacterium* were abundant was slightly enriched in iron, manganese and oxygen (Table 1 and Figure 6). This agrees with the observation that greater abundance of iron reducers and oxidisers were predicted in the distal and proximal wells, respectively (Figure 8). Similarly, the greater abundance of iron-reducers in the background well can be attributed to *Rhodoferax*, although iron in this well was below the limit of detection. Like sulphate reduction, iron reduction must be limited due to the low organic matter.

#### Other functions

Genus *Polynucleobacter* was one of the abundant taxa (Figure 4). Members of the genus are obligate endosymbionts of ciliates (Heckmann and Schmidt, 1987), and thus the presence of a substantial protistan population is indicated, which is further supported by the presence of Legionella, a parasite of amoebae. In addition, *Polynucleobacter*, and other genera such as *Aquabacterium*, *Rhodoferax*, *Duganella* and *Limnohabitans*, were disproportionately more abundant in spring (Figure S7). The seasonal variation in the microbial community composition will be addressed in a feature manuscript (in preparation). *Thermomonas* and *Sediminibacterium* were also among the abundant OTUs, but none has documented bioremediation/biodegradation potential. *Undibacterium* and *Duganella* equally formed part of the abundant OTUs. Culture-based identification revealed that *Undibacterium* composed of *U. parvum* and *U. pigrum*, both freshwater flora, while *Duganella* was not detected by the culture method, but showed higher abundance in the background and distal wells (Figure 4). The genus showed stronger correlation with dissolved oxygen (Figure 6) and given that dissolved oxygen was higher in these wells, it might be a relevant selection factor for the genus.

## Conclusion

This study reports the microbial diversity of a landfill-contaminated confined aquifer surveyed in spring and autumn, 2018. Both culture-dependent and culture-independent microbial techniques were applied. Comparison of microscopic cell counts and plate counts as well as metabarcoding data suggest that the microbes in the most contaminated parts of the aquifer were mostly culturable heterotrophs. Comparison between uncontaminated and contaminated aquifer showed higher microbial diversity in the contaminated aquifer. Within the contaminated aquifer, microbial diversity was moderate in the proximal well, highest in the intermediate well and lowest in the distal well. The lower diversity in the distal well indicates dominance of the microbial community by a few taxa. Each well had a distinctive microbial flora, but the proximal and intermediate wells seemed to be ecologically related as they had more similar microbial community composition. There was a clear difference between the flora of the contaminated wells and the uncontaminated well. The data suggests that the microbiome of the contaminated aquifer has been impacted by the landfill leachate. Functional analysis indicates the presence of microbes capable of hydrocarbon, sulphur, nitrogen, iron and manganese metabolism.

## Acknowledgement

We thank Frode Bergan and Tom Aage Aarnes for participation in fieldworks, Karin Brekke Li for technical assistance in water chemistry analysis and Benedikte N. Pedersen (PhD) for technical advice during sequencing of bacterial isolates. Fluorescence microscopy was conducted at the department of process, energy and environment, University of South-Eastern Norway and we thank Professor Rune Bakke (PhD) for providing access to the instrument and Ikumi Umetani (PhD) for help with its use. The sequencing service was provided by the Norwegian Sequencing Centre (https://www.sequencing.uio.no/), a national technology platform hosted by the University of Oslo and supported by the Functional Genomics and Infrastructure programs of the Research Council of Norway and the Southeastern Regional Health Authorities.

## Supporting information

### Methods

Method S1.

Method S2.

### Figures

Figure S1.

Figure S2.

Figure S3.

Figure S4.

Figure S5.

Figure S6.

Figure S7.

### Tables

Table S1.

Table S2.

Table S3.

Table S4.

Table S5.

